# Selective Nigrostriatal Dopamine Excess Impairs Behaviors Linked to the Cognitive and Negative Symptoms of Psychosis

**DOI:** 10.1101/2022.08.09.503421

**Authors:** Nicolette A. Moya, Seongsik Yun, Stefan W. Fleps, Madison M. Martin, Jacob A. Nadel, Lisa R. Beutler, Larry S. Zweifel, Jones G. Parker

## Abstract

**BACKGROUND:** Excess dopamine release in the dorsal striatum (DS) is linked to psychosis. Antipsychotics are thought to work for positive symptoms by blocking striatal D2 dopamine receptors, but they lack efficacy for the negative and cognitive symptoms. Further, broadly increasing dopamine release improves cognitive function. These observations fueled the dogma that excess dopamine is not involved in negative and cognitive symptoms, but this has never been tested with dopamine pathway specificity.

**METHODS:** We selectively re-expressed excitatory TRPV1 receptors in DS-projecting dopamine neurons of male and female *Trpv1* knockout mice. We treated these mice with capsaicin (TRPV1 agonist) to selectively activate these neurons, validated this approach with fiber photometry, and assessed its effects on social and cognitive function. We combined this manipulation with antipsychotic treatment (haloperidol) and compared the pathway-specific manipulation to treatment with the non-selective dopamine releaser amphetamine.

**RESULTS:** Selectively activating DS-projecting dopamine neurons increased DS (but not cortical) dopamine release and increased locomotor activity. Surprisingly, this manipulation also impaired behavioral processes linked to negative and cognitive symptoms (social drive and working memory). Haloperidol normalized locomotion, only partially rescued working memory, and had no effect on social interaction. By contrast, amphetamine increased locomotion but did not impair social interaction or working memory.

**CONCLUSIONS:** Excess dopamine release, when restricted to the DS, causes behavioral deficits linked to negative and cognitive symptoms. Previous studies using non-selective approaches to release dopamine likely overlooked these contributions of excess dopamine to psychosis. Future therapies should address this disregarded role for excess striatal dopamine in the treatment-resistant symptoms of psychosis.

## INTRODUCTION

Current antipsychotic treatments are largely ineffective for the cognitive and negative symptoms of psychosis (1). Negative symptoms such as the loss of desire for social engagement and cognitive symptoms like deficits in working memory are just as debilitating to quality of life as the hallmark positive symptoms of psychosis (*e*.*g*., hallucinations and delusions) (2). Therefore, there is an immediate need to address these treatment-resistant symptoms. A major barrier to this has been an imprecise understanding of the underlying neural substrates of psychosis and how its neuropathology maps onto different symptoms.

For decades, we have known that the neurotransmitter dopamine plays an important role in psychosis, but the mechanisms underlying this role remain unclear. Nearly all antipsychotic drugs block D2 dopamine receptors (D2Rs), and functional imaging studies show that schizophrenia patients have increased dopamine in the DS, where D2Rs are abundant (3-6). These observations fueled the prominent hypothesis that excess striatal dopamine is responsible for the D2R antagonist-responsive, positive symptoms of schizophrenia but not its D2R antagonist-resistant, negative and cognitive symptoms (7). This idea is further supported by the fact that amphetamine (a dopamine releasing drug) can acutely induce positive symptoms (8) but actually improves cognitive function in schizophrenia patients (9).

However, D2Rs are not the only dopamine receptors in the striatum, which equally expresses D1 dopamine receptors (D1Rs) (10). Because D1Rs are not a principal target of existing antipsychotics, their signaling may contribute to symptoms that are resistant to treatment with D2R antagonist-based antipsychotics. Moreover, the pro-cognitive effects of amphetamine in schizophrenia patients could stem from the fact that it increases dopamine release throughout the brain (not only in the dorsal striatum, as observed in schizophrenia), including in the prefrontal cortex (PFC) where dopamine release is actually decreased in these patients (10-12). Moreover, the dorsal striatum is anatomically interconnected with the PFC and other brain structures implicated in cognitive and social function (13-15). These circuit-level interactions and the specificity of antipsychotics for D2Rs suggest that excess dopamine signaling, when restricted to the DS, could contribute to the negative and cognitive symptoms of psychosis. This idea that has never been directly tested.

To explore this, we developed an experimental approach using the capsaicin-gated ion channel, TRPV1, to accurately recapitulate the pathway-specific increase in dopamine transmission observed in schizophrenia and examined its effects on antipsychotic treatment-responsive and resistant behavioral processes in mice. In contrast to the prevailing view, selectively driving DS dopamine transmission impaired behavioral processes related to the negative and cognitive symptoms of psychosis (social interaction and working memory). These behavioral deficits were largely unresponsive to the D2R-antagonist/antipsychotic drug haloperidol and did not occur following treatment with the non-selective dopamine releaser amphetamine. Our findings suggest that excess striatal dopamine plays a broader role in the symptomatology of psychosis than previously thought. This insight exposes a gap in our understanding of dopamine’s role in psychosis and the tools established here provide a path forward to close this gap and develop more comprehensive antipsychotic treatments.

## METHODS AND MATERIALS

### Animals

We housed and handled all mice according to guidelines approved by the Northwestern University Animal Care and Use Committee. We used both male and female mice housed on a reverse light cycle for all experiments. For capsaicin experiments, we crossed homozygous DAT^IREScre^ mice (Jax #0006660) with *Trpv1* knockout (KO) mice (Jax #003770) to generate double heterozygous mice that we then crossed to generate DAT^cre/+^; TRPV1 KO mice (16, 17). We maintained all founder mouse lines through backcrossing to C57BL/6J mice (Jax #000664), the same mouse strain we used for behavioral experiments with amphetamine. All mice were 12–24 weeks at the start of experimental testing. Detailed materials and methods are provided in the **Supplementary Methods**.

### Drugs

We injected all drugs subcutaneously at volume of 10 mL·kg^-1^. We dissolved capsaicin (3.5–10 mg·kg^-1^; Alomone Labs) in 3.33% Tween 80 in PBS, D-Amphetamine hemisulfate (0.5–10 mg·kg^-1^; Sigma) in saline, and haloperidol (0.032–0.1 mg·kg^-1^; Sigma) in 0.3% tartaric acid.

### Surgical Procedures

We stereotaxically injected either AAV2/5-hSyn-FLEX-TRPV1-mCherry or AAV2/5-hSyn-DIO-mCherry bilaterally in the substantia nigra pars compacta (SNc) of DAT^cre/+^; TRPV1 KO mice. For animals used for fiber photometry, we also injected AAV2/9-CAG-dLight1.3b into the dorsomedial striatum (DMS) and medial prefrontal cortex (mPFC; unilaterally), followed by fiberoptic cannula implants into the same regions (**Figure 1A, 2A**).

**Figure 1.**
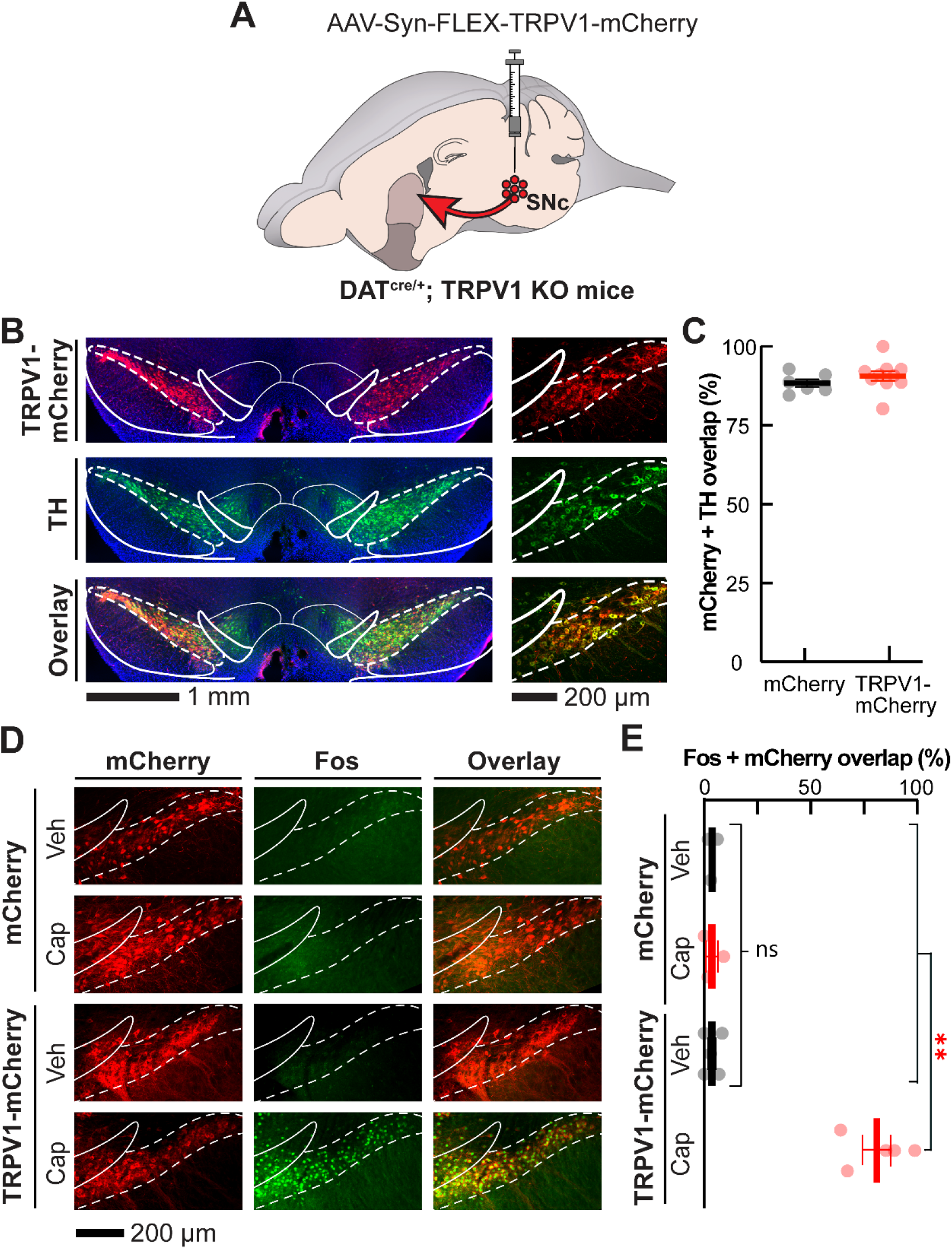
Histological validation of the selective expression of functional TRPV1 in SNc dopamine neurons. **A)** We virally expressed Cre-dependent mCherry (control) or TRPV1-mCherry (experimental) in DS-projecting SNc dopamine neurons of DAT^cre/+^; TRPV1 KO mice. **B)** Coronal brain sections containing SNc and VTA from a representative experimental (TRPV1-mCherry) mouse shown at 10× (*left*) and 16× resolution (*right*; *blue*: DAPI nuclear stain; *red*: α-RFP; *green*: α-TH). **C)** Mean ± s.e.m. percentage of mCherry-expressing neurons that are TH positive in control and experimental mice (*N* = 6 and 10 respectively; average of 3 brain slices per mouse). **D)** Coronal brain sections containing SNc from representative control (*top*) and experimental (*bottom*) mice immunostained for Fos (*green*) or RFP (*red*) 75 min after systemic vehicle or capsaicin treatment (10 mg·kg^-1^). **E)** Mean ± s.e.m. percentage of mCherry-expressing neurons that are Fos positive as a function of treatment and experimental group (*N* = 3–5 mice per group; average of 3 brain slices per mouse). In all images, white dashed lines indicate boundaries of SNc and solid lines indicate boundaries of adjacent brain structures. \*\**P* < 0.01 comparing vehicle to capsaicin treatment; Unpaired t-test in **C**; Kruskal-Wallis unpaired one-way ANOVA in **E**. Details for these and all other statistical comparisons are presented in the **Supplemental Table**.

**Figure 2.**
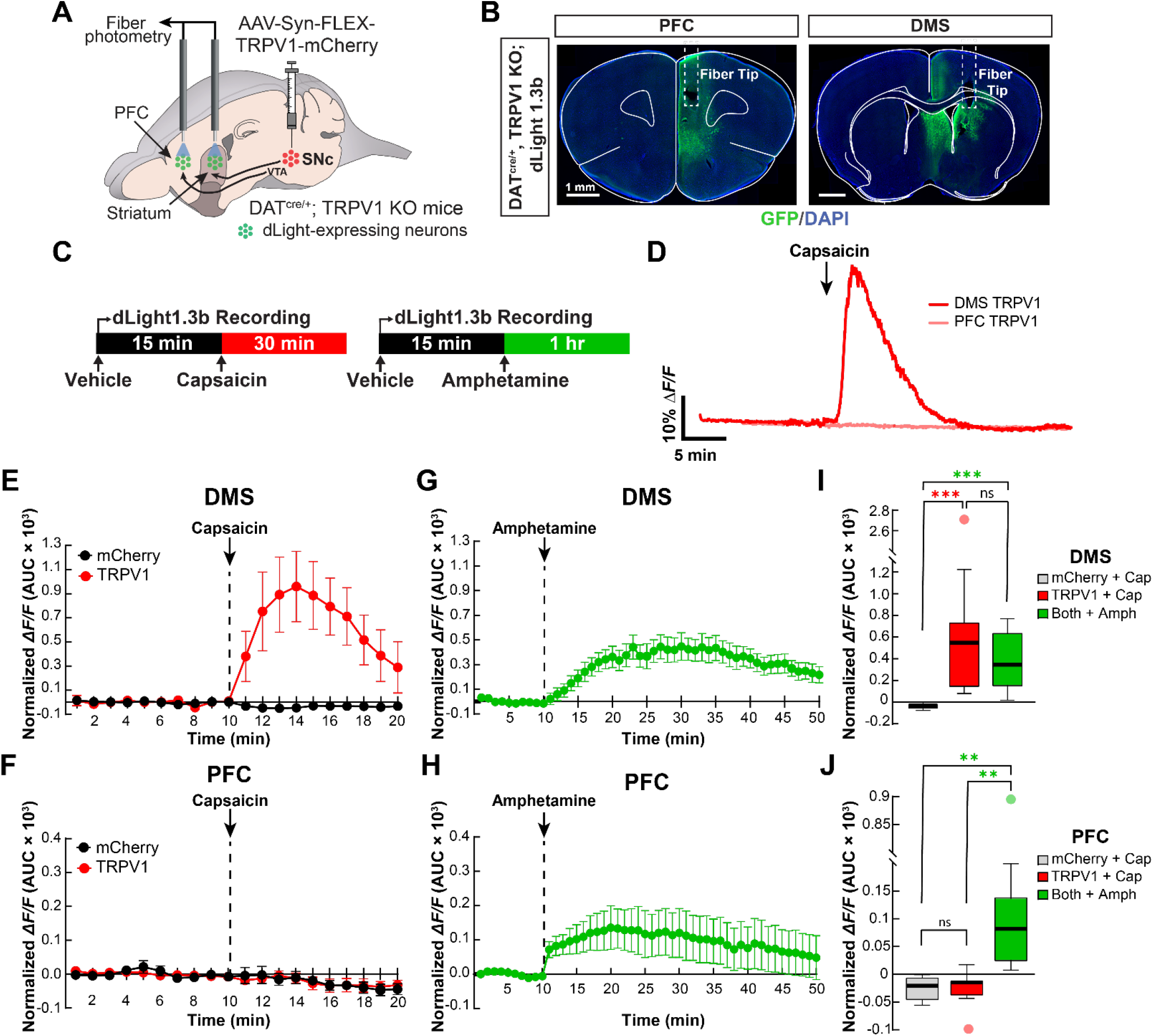
Recordings of dLight1.3b fluorescence in mPFC and DMS following systemic capsaicin or amphetamine treatment. **A)** We virally expressed mCherry (*control*) or TRPV1-mCherry (*experimental*) in DS-projecting SNc dopamine neurons of DAT^cre/+^; TRPV1 KO mice and the fluorescent dopamine sensor dLight1.3b in mPFC and DMS. We then implanted fiber-optic cannulas and used fiber photometry to simultaneously record dopamine transmission in mPFC and DMS. **B)** Representative coronal brain sections from an experimental, TRPV1 mouse expressing dLight1.3b in prefrontal cortex and dorsomedial striatum (*green*: α -GFP; *blue*: DAPI nuclear stain; scale bar: 1 mm). White dashed lines indicate the position of the implanted fiber-optic probe. **C)** Schematic representation of the time course of dLight1.3b recordings and drug treatments. We administered vehicle at t = 0 min, and capsaicin (10 mg·kg^-1^) or amphetamine (10 mg·kg^-1^) at t = 15 min. We recorded dLight1.3b fluorescence for 15-, 30-, or 60-min following vehicle, capsaicin, or amphetamine treatment, respectively. **D)** Example trace (% *ΔF/F*) from of dLight1.3b fluorescence in mPFC and DMS in response to capsaicin treatment in a representative TRPV1 mouse. **E, F)** Mean ± s.e.m. dLight1.3b fluorescence in DMS, **E**, and mPFC, **F**, calculated using area-under-the-curve (AUC) in 1-min time bins and normalized to values in the final 10 min following vehicle treatment in control (mCherry) and experimental (TRPV1) mice. Capsaicin responses are truncated to the first 10 min when changes in dopamine were most evident. **G, H)** Mean ± s.e.m. dLight1.3b fluorescence in DMS, **G**, and mPFC, **H**, calculated using AUC in 1-min time bins and normalized to values in the final 10-min following vehicle treatment in control and TRPV1 mice. Amphetamine responses are truncated to the first 40 min when changes in dopamine were most evident. **I, J)** dLight1.3b fluorescence in DMS, **I**, and mPFC, **J**, calculated using AUC in 1-min time bins, normalized to values in the final 10-min following vehicle treatment, and averaged across the 10, 1-min time bins surrounding the peak response following systemic capsaicin or amphetamine treatment in control and TRPV1 mice. Both control and experimental groups were combined in the analysis of amphetamine treatment effects (**G**–**J**). In these and all other box-and-whisker plots, the horizontal lines denote median values, boxes cover the middle two quartiles and whiskers span 1.5× the interquartile range. *N* = 7 mCherry, *N* = 11 TRPV1, and *N* = 18 combined amphetamine mice; \*\*\**P* < 0.001 and \*\**P* < 0.01; Holm-Sidak’s multiple comparison test.

### Fiber Photometry

We used a commercial fiber photometry system and software (TDT) to record each mouse’s baseline dLight1.3b activity in the mPFC and DMS for 15 min, subcutaneously injected vehicle, recorded for an additional 15 min, and then subcutaneously injected capsaicin or amphetamine (10 mg·kg^-1^ each) and recorded dLight1.3b activity for 30 or 60 min, respectively (**Figure 2C**). We used custom MATLAB scripts to compute normalized dLight1.3b fluorescent traces (% Δ*F/F*) and area-under-the-curve (AUC) measurements for traces in 1-min time bins.

### Open Field Locomotor Activity

We habituated mice to the open field for 20 min, injected vehicle, recorded locomotion for 30 min or 1 h, injected capsaicin or amphetamine at varying doses (one dose per day) and recorded locomotion for 30 min or 1 h, respectively (**Figure 3B, 5A**). For haloperidol experiments, we pretreated mice with varying doses of haloperidol (one dose per day), recorded locomotion for 30 min, injected capsaicin (3.5 mg·kg^-1^) and again recorded locomotion for 30 min (**Figure 6A**). To analyze locomotor speed, we used a video camera and custom software written in ImageJ to track each mouse’s position over time in the open field (18). The open field locomotor experiments for mice treated with vehicle followed by capsaicin (3.5 mg·kg^-1^) were experimentally equivalent and the data statistically indistinguishable between the first cohort of mice (**Figure 3B, C**) and the cohort used for the haloperidol experiments (**Figures 6A, B**), so we combined all mice in those treatment groups for statistical comparisons.

**Figure 3.**
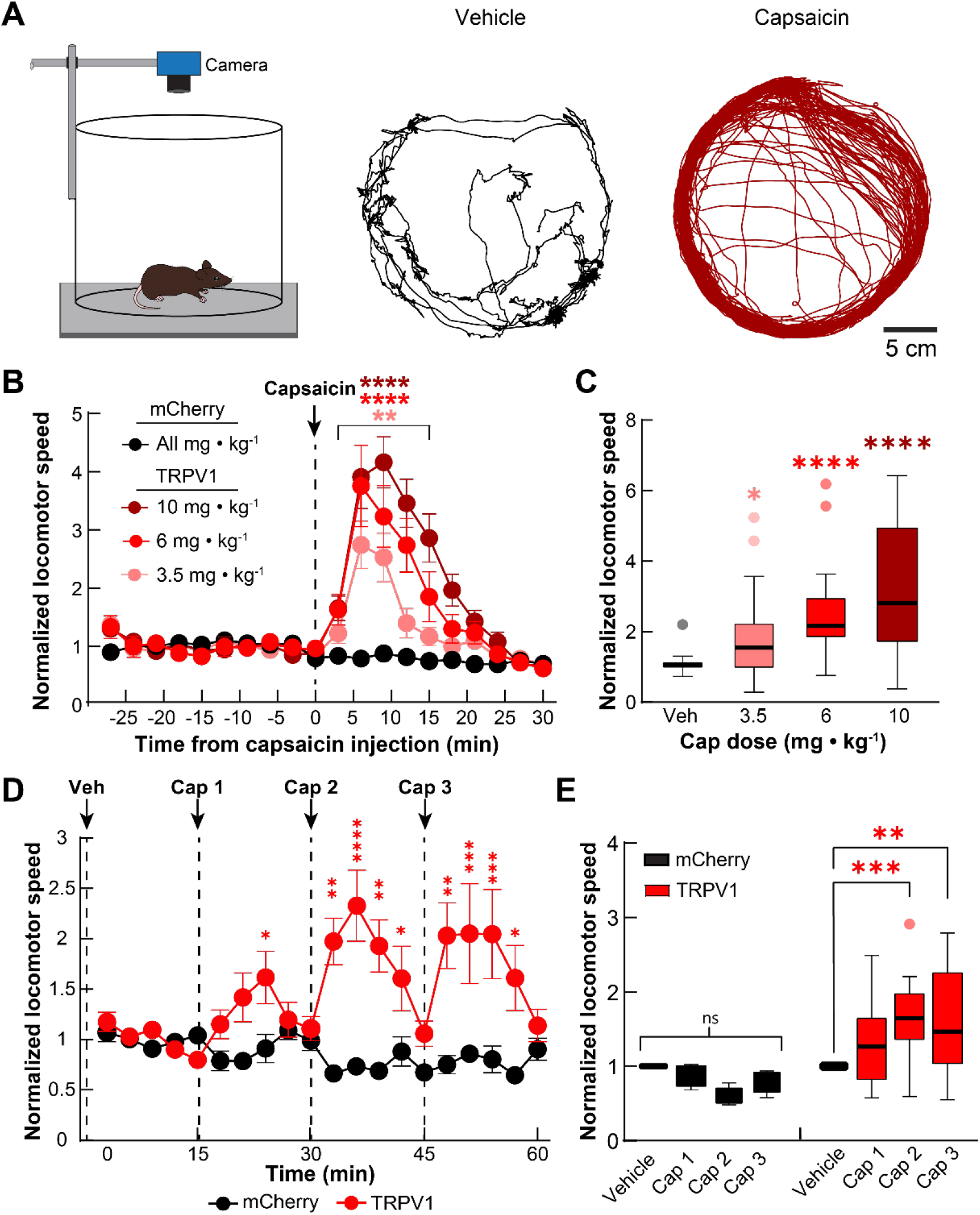
Selective nigrostriatal dopamine transmission dose-dependently increases locomotion in experimental TRPV1, but not control mice. **A)** We recorded locomotor activity in an open field arena for 30-min following vehicle then capsaicin treatment (*left*). Example trajectories (10-min duration) for an experimental (TRPV1) mouse following systemic treatment with vehicle (*middle*) or capsaicin (*right*; 10 mg·kg^-1^). **B, C)** Systemic capsaicin treatment dose-dependently increased locomotor speed in experimental but not control mice. Data in **B** are mean ± s.e.m. normalized to values following vehicle treatment and plotted in 3-min time bins; the control data are averaged across all capsaicin doses. Data in **C** are averaged during the first 15-min following vehicle or capsaicin treatment (*N* = 8 mCherry and *N* = 29–30 TRPV1 mice). **D, E)** Locomotor response to repeated capsaicin injections (3.5 mg·kg^-1^ every 15 min) in experimental and control mice. Data in **D** are mean ± s.e.m. normalized to values following vehicle treatment (t = 0–15 min) and plotted in 3-min time bins. Data in **E** are 15-min averages following each injection, normalized to associated values following vehicle treatment (*N* = 5 mCherry and *N* = 10 TRPV1 mice). \*\*\*\**P* < 0.0001, \*\*\**P* < 0.001, \*\**P* < 0.01 and \**P* < 0.05 comparing capsaicin and vehicle treatments; two-way ANOVA in **B**; Holm-Sidak’s multiple comparison test in **C**–**E**.

### Juvenile Social Exploration

We habituated individual mice in their home cage for 10 min, injected vehicle, capsaicin (3.5 mg·kg^-1^), or amphetamine (1 mg·kg^-1^), waited 2 min (capsaicin) or 4 min (amphetamine), and placed a novel juvenile (postnatal day 21–35) conspecific mouse in the adult’s home cage (**Figure 4A**). For haloperidol experiments, we pretreated mice with either vehicle or haloperidol (0.1 mg·kg^-1^), waited 10 min, then injected capsaicin (3.5 mg·kg^-1^), waited 2 min, and placed the novel juvenile in the adult’s home cage (**Figure 6C**). On each day, we measured the amount of time the adult experimental mouse spent interacting with the juvenile test subject for 5 min, which included the sum duration of sniffing, grooming, approaching, or pawing initiated by the adult mouse towards the juvenile (19).

**Figure 4.**
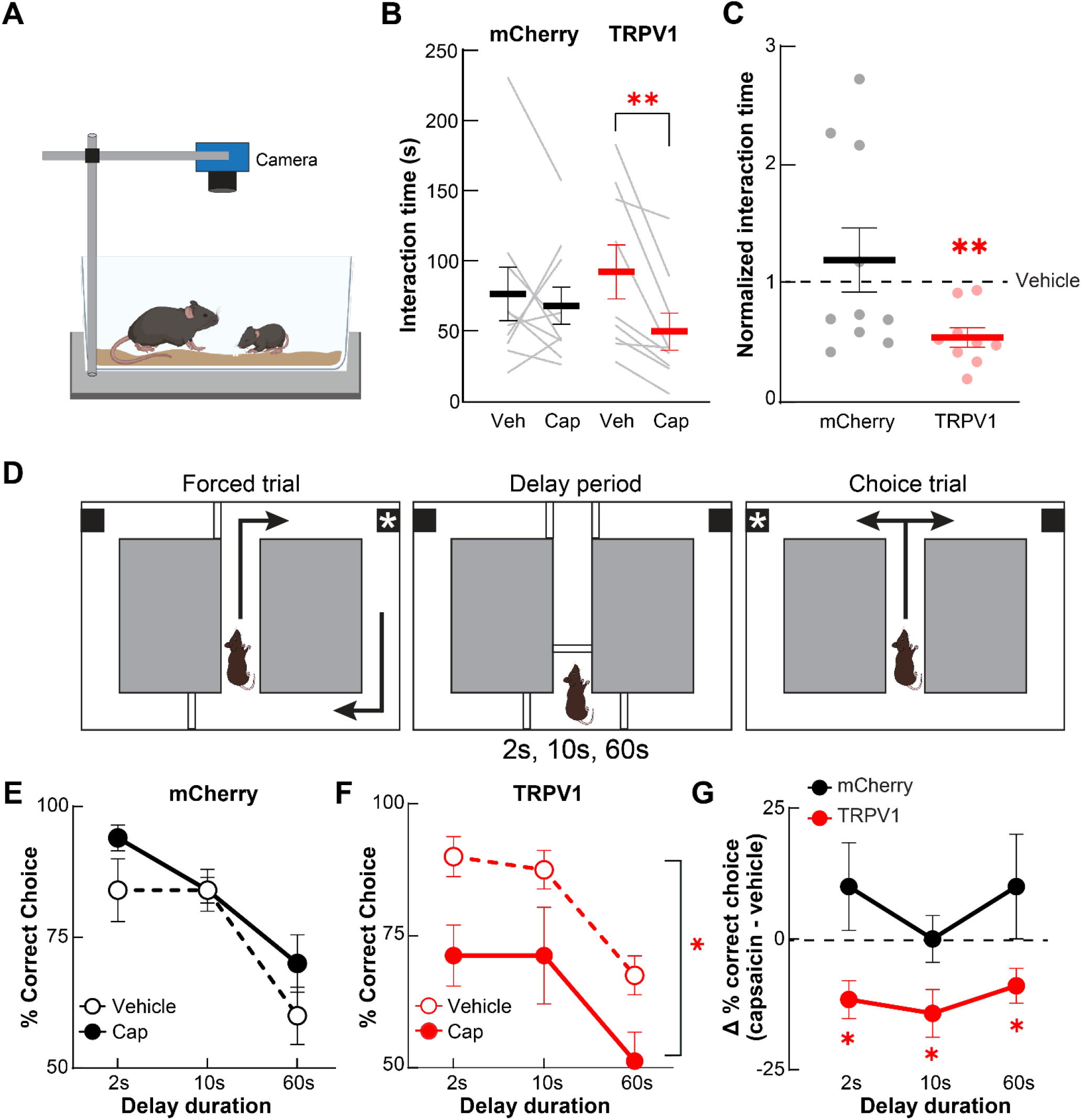
Selective nigrostriatal dopamine excess disrupts social and cognitive function. **A)** We recorded behavior during an assay of juvenile social exploration in which control (mCherry) and experimental (TRPV1) adult mice interacted with a juvenile, conspecific mouse following vehicle or capsaicin (3.5 mg·kg^-1^) treatment for 5 min. **B, C)** Mean ± s.e.m. interaction time, quantified as the sum duration of sniffing, grooming, approaching, or pawing initiated by the adult mouse towards the juvenile, **B**, or normalized to values following vehicle treatment, **C**, decreased following capsaicin treatment in experimental, but not control mice (*N* = 10 mCherry and *N* = 9 TRPV1 mice). **D)** We used an automated T-maze to evaluate spatial working memory with a delayed non-match to sample (DNMS) protocol, where mice alternated between forced and choice trials with varying delay periods (2, 10 or 60 s) in a center holding area between trials. **E, F)** Mean ± s.e.m. percent correct choice in control, **E**, and experimental, **F**, mice following vehicle or capsaicin treatment (3.5 mg·kg^-1^) in the DNMS task at different delay periods. **G)** Mean ± s.e.m. difference in percent correct choice following capsaicin and vehicle treatment in experimental and control mice (*N* = 5 mCherry and *N* = 8 TRPV1 mice; \*\**P* < 0.01 and \**P* < 0.05 comparing capsaicin and vehicle treatments; Wilcoxon signed-rank test in **B, C**; two-way ANOVA in **E, F**; Holm-Sidak’s multiple comparison test in **G**).

### Spatial Working Memory

We individually housed and gradually food restricted mice to 85% of their *ad libitum* body weight. We then habituated mice to an automated T-maze (Maze Engineers) for 2 days. Following habituation, we pre-trained mice in sessions consisting of 10 alternating (left vs. right) forced trials from the center holding area to a baited goal arm and then back to the center holding area. The day after pre-training, we trained mice in a delayed non-match to sample (DNMS) task in which we interleaved forced and choice trials (20). During the choice trial, mice were required to choose the arm opposite to the immediately preceding forced trial to obtain a food reward. Once mice had at least three consecutive days with ≥70% correct choice trials in DNMS training, we proceeded with pharmacological testing. In each session, we injected mice with vehicle, capsaicin (3.5 mg·kg^-1^), or amphetamine (1 mg·kg^-1^), waited 2 min (capsaicin) or 4 min (amphetamine), and tested their DNMS performance with a delay period of 2, 10, or 60 s between forced and choice trials on sequential days (**Figure 4E–G, 5F–G**). For sessions that exceeded 15 min, we administered additional capsaicin injections every 15 min to sustain the drug’s behavioral effect (**Figure 3B**). For haloperidol experiments, we pretreated mice with either vehicle or haloperidol (0.1 mg·kg^-1^), waited 10 min, then injected capsaicin (3.5 mg·kg^-1^), waited 2 min, and tested their performance with the same 2, 10, and 60 s delays while administering capsaicin every 15 min until task completion (**Figure 6E– G**).

### Histology

Following all photometry and behavioral experiments, we euthanized and intracardially perfused mice, then removed and sliced their brains for immunostaining. We used α-TH or α-Fos with α-RFP antibodies to quantify TH or Fos colocalization with mCherry in SNc. For dlight1.3b analysis, we used an α-GFP antibody in DMS and mPFC. We imaged fluorescence using a multiphoton or wide-field fluorescence microscope to verify the accuracy of injection and/or implantation (**Figure 1, 2B**).

### Statistical Analysis

We used Prism (GraphPad) to perform all statistical tests. For comparisons between data that did not conform to a normal distribution, we used non-parametric statistical comparisons. For comparisons of more than two groups, we used one-way ANOVA. For comparisons of two or more groups across conditions or time, we used two-way repeated measures ANOVA. For all *post-hoc* analyses, we used Holm-Sidak’s correction for multiple comparisons. All statistical comparisons presented are in the **Supplementary Table**.

## RESULTS

### Selective Expression and Activation of TRPV1 in SNc Dopamine Neurons

To selectively drive dopamine release in the DS, we re-expressed the capsaicin-sensitive, excitatory cation channel TRPV1 in DS-projecting, substantia nigra pars compacta (SNc) dopamine neurons. Specifically, we bilaterally injected a Cre-dependent virus expressing TRPV1-mCherry (or mCherry control) into the SNc of DAT^cre/+^ mice that lack the endogenous *Trpv1* gene (DAT^cre/+^; TRPV1 KO mice) (**Figure 1A**). This approach resulted in a high degree of overlap between neurons that express mCherry and neurons that were immunopositive for the dopamine biosynthesis enzyme tyrosine hydroxylase (TH), specifically in the SNc (**Figure 1B, C**). Considering these mice lack the *Trpv1* gene, this allowed us to inject them systemically with the TRPV1 agonist capsaicin and selectively activate neurons at the site of virus injection without engaging endogenous TRPV1 receptors in the periphery (21, 22). To confirm that virally expressed TRPV1 was functional in SNc dopamine neurons, we systemically administered vehicle or capsaicin (10 mg·kg^-1^) to control and experimental (TRPV1) mice. 75-min following injection, there was a high degree of overlap between neurons expressing mCherry and the immediate early gene product Fos in experimental mice after capsaicin treatment. By contrast, there was little to no overlap of Fos with mCherry in control mice injected with capsaicin, or in TRPV1 mice injected with vehicle (**Figure 1D, E**). These results indicate that TRPV1 expression was restricted to SNc dopamine neurons in these mice and that systemic capsaicin treatment selectively activated these neurons.

### Selective Activation of Nigrostriatal Dopamine Release In Vivo

To confirm that capsaicin treatment increases dopamine release in a pathway-specific manner in these mice, we virally expressed the fluorescent dopamine sensor dLight (AAV2/9-CAG-dLight1.3b) and implanted fiberoptic probes unilaterally into the dorsomedial striatum (DMS) and medial prefrontal cortex (mPFC) for dual-site, fiber photometry recordings (**Figure 2A, B**). We then recorded dLight fluorescence in the mPFC and DMS following systemic treatment with vehicle, capsaicin, or the non-selective dopamine releaser amphetamine (**Figure 2C**). Capsaicin treatment (10 mg·kg^-1^) increased dopamine transmission (Δ*F/F*) in the DMS of TRPV1, but not control mice (**Figure 2D, E**). Importantly, capsaicin treatment did not increase dopamine release in the mPFC of either TRPV1 or control mice (**Figure 2D, F**).

In contrast to capsaicin, amphetamine treatment (10 mg·kg^-1^) increased dopamine release in both the DMS and mPFC of these mice (**Figure 2G, H**), indicating that dLight1.3b is suitable for measuring dopamine release, even in the mPFC, where dopamine levels are lower. Taken together, these findings demonstrate that our approach selectively drives activity in DS-projecting, SNc dopamine neurons and mimics the pathway-specific excess of dopamine observed in schizophrenia patients (12) (**Figure 2I, J**).

### Selective Nigrostriatal Dopamine Excess Increases Locomotor Activity

Locomotor hyperactivity is associated with the positive symptoms of psychosis in that both are reversed by effective antipsychotic treatments (23). We recorded the locomotor activity of control and TRPV1 mice in an open field arena following treatment with a range of capsaicin doses (**Figure 3A**). Capsaicin treatment dose-dependently induced locomotor hyperactivity in TRPV1, but not control mice (**Figure 3B, C**). The effect of capsaicin on locomotion started rapidly (< ∼3 min) and lasted about 15 min after injection for the lowest dosage tested (3.5 mg·kg^-1^; **Figure 3B**). To determine whether we could repeatedly inject capsaicin without de-sensitizing its behavioral effects, we administered three successive injections separated by 15 min. These repeated capsaicin treatments consistently induced hyperlocomotion in TRPV1 mice, but not control mice, with no evidence of desensitization (**Figure 3D, E**). These results show that selectively driving DS dopamine release with capsaicin increases locomotor activity and establishes the time-course and dose-dependence of this effect. Given that the lowest dose tested here (3.5 mg·kg^-1^) only modestly increased locomotion and reliably elicited a behavioral response with no obvious desensitization, we used this dose for all subsequent behavioral assessments.

### Selective Nigrostriatal Dopamine Excess Disrupts Social and Cognitive Function

The negative and cognitive symptoms of psychosis include social withdrawal and deficits in working memory. To determine if selectively driving DS dopamine transmission affects behaviors associated with these symptoms, we evaluated the effects of capsaicin treatment on juvenile social exploration (JSE) and spatial working memory (WM). Surprisingly, capsaicin treatment reduced social interaction with a juvenile conspecific mouse in TRPV1, but not control mice (**Figure 4A–C**). Likewise, capsaicin treatment disrupted working memory in TRPV1 but not control mice, in a T-maze, delayed non-match to sample (DNMS) task requiring mice to choose the arm opposite from the one visited on the preceding forced-choice trial (**Figure 4D–G**). Notably, capsaicin treatment did not affect performance in a version of the T-Maze task requiring mice to repeatedly choose the same reward arm where working memory was not required (mean ± s.e.m: 79 ± 2 and 75 ± 5 % correct choice following vehicle and capsaicin treatment, respectively; *N* = 4 TRPV1 mice; *P* = 0.75; Wilcoxon signed-rank test). These results indicate that excess dopamine, when restricted to the DS, diminishes cognitive and social function.

### Non-Selective Dopamine Excess Alters Locomotor Activity but not Social or Cognitive Function

Amphetamine is a psychostimulant that is often used to approximate dopamine dysregulation in psychosis in rodents (23). In contrast to the viral-genetic approach used here, amphetamine increases dopamine release throughout the brain (not selectively in DS), including in the PFC (24). Consistent with previous studies, treating C57BL/6J mice with amphetamine dose-dependently increased locomotor activity in the open field (**Figure 5A, B**). A 1 mg·kg^-1^ amphetamine dose increased locomotor speed most comparably to the 3.5 mg·kg^-1^ dose of capsaicin we used for the WM and JSE experiments in TRPV1 mice (**Figure 5C**), so we used this dose for subsequent behavioral assays. Despite inducing similar levels of locomotion, amphetamine treatment had no effect on social interaction or working memory performance compared to vehicle treatment (**Figure 5D–G**). These results suggest that the behavioral deficits caused by capsaicin treatment in TRPV1 mice were not solely due to changes in their locomotor activity. Moreover, these findings highlight the differences between the effects of DS-restricted and brain wide excess in dopamine, in that only the former produces deficits in behaviors related to the negative and cognitive symptoms of psychosis (**Figures 2, 4, 5**).

**Figure 5.**
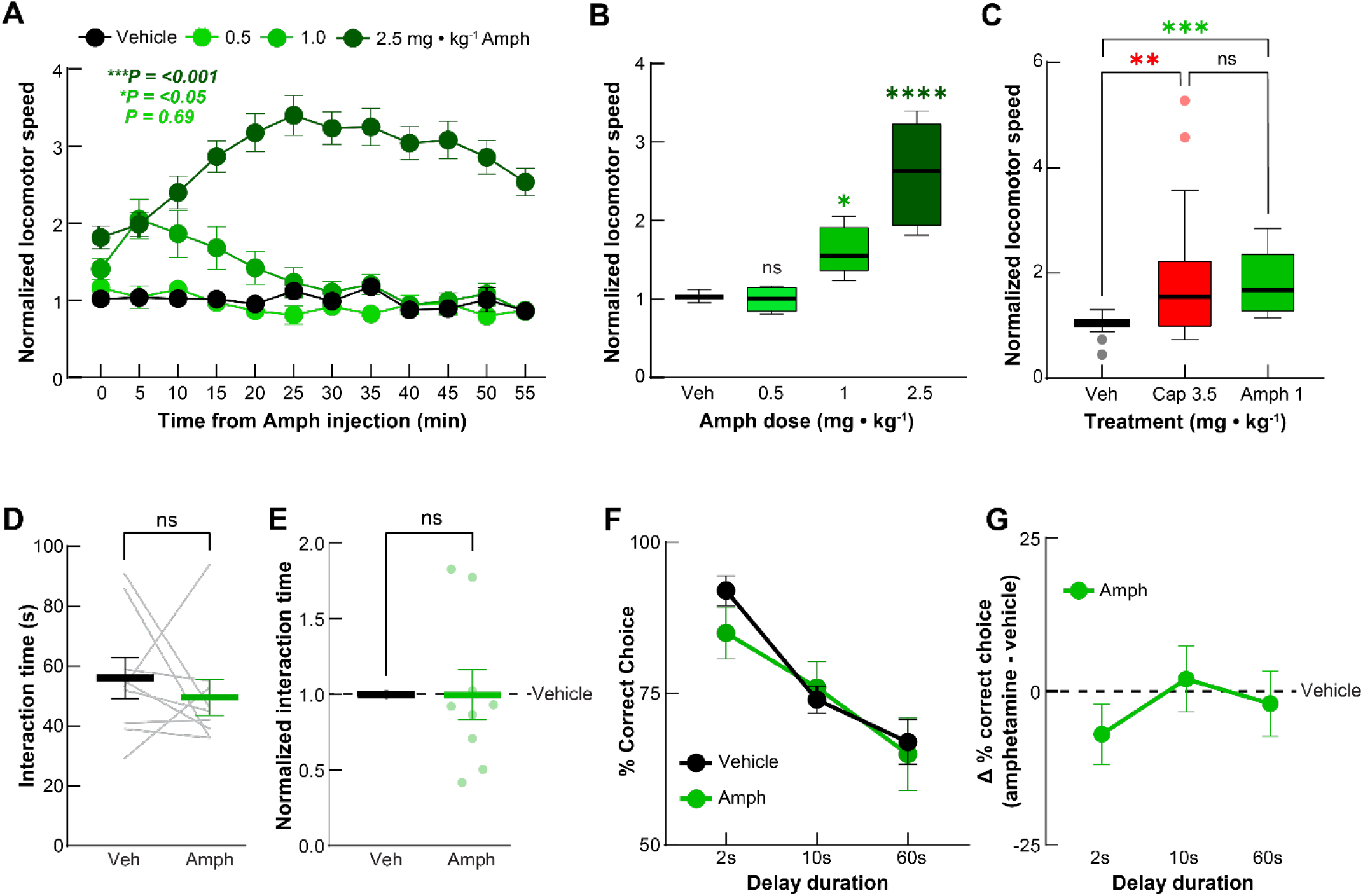
Amphetamine treatment induces hyperlocomotion but has no effect on social exploration or working memory. **A, B)** Systemically treating C57BL/6J mice with amphetamine dose-dependently increased locomotor speed in the open field. Data in **A** are mean ± s.e.m. normalized to vehicle treatment and plotted in 5-min time bins. Data in **B** are 15-min averages following each injection, normalized to values following vehicle treatment. **C)** Amphetamine treatment (1 mg·kg^-1^) in C57BL/6J mice induced locomotion equivalently to capsaicin treatment (3.5 mg·kg^-1^) in TRPV1 mice in the 15-min following injection, normalized to values following vehicle treatment. **D, E)** Amphetamine treatment (1 mg·kg^-1^) did not significantly affect the time C57BL/6J mice spend interacting with a juvenile conspecific, **D**, and there was no difference, **E**, in social interaction time when normalized, within each mouse, to values following vehicle treatment (mean ± s.e.m.). **F, G)** Amphetamine treatment (1 mg·kg^-1^) did not significantly affect the percent of correct choices C57BL/6J mice made in the T-maze, DNMS task, **F**, and there was no difference, **G**, in the percentage of correct choices when normalized, within each mouse, to values following vehicle treatment (mean ± s.e.m.; *N* = 9–10 C57BL/6J mice and *N* = 29 TRPV1 mice; \*\*\*\**P* < 0.0001, \*\*\**P* < 0.001, and \*\**P* < 0.01 comparing amphetamine, capsaicin, and vehicle treatments; Holm-Sidak’s multiple comparison test in **A**–**C, F, G**; Wilcoxon signed-rank test in **D, E**).

### Haloperidol Normalizes Locomotor Hyperactivity, and Partially Normalizes Cognitive but not Social Deficits Caused by Selective Nigrostriatal Dopamine Excess

Classical antipsychotic drugs are largely ineffective for treating the negative and cognitive symptoms of psychosis (1). To determine whether the common antipsychotic drug haloperidol normalizes the behavioral deficits of a selective excess in DS dopamine, we tested its effects in TRPV1 mice. Pretreating these mice with a range of haloperidol doses blocked their increase in locomotor activity following systemic capsaicin treatment (**Figure 6A, B**). For subsequent behavioral tests in TRPV1 mice, we used the highest dose of haloperidol tested in the open field (0.1 mg·kg^-1^). Despite normalizing capsaicin-induced locomotion, haloperidol pretreatment failed to normalize the capsaicin-induced decrease in social interaction in TRPV1 mice (**Figure 6C, D**). Furthermore, haloperidol only partially rescued the disruption of working memory caused by capsaicin treatment in these mice (**Figure 6E, F**).

**Figure 6.**
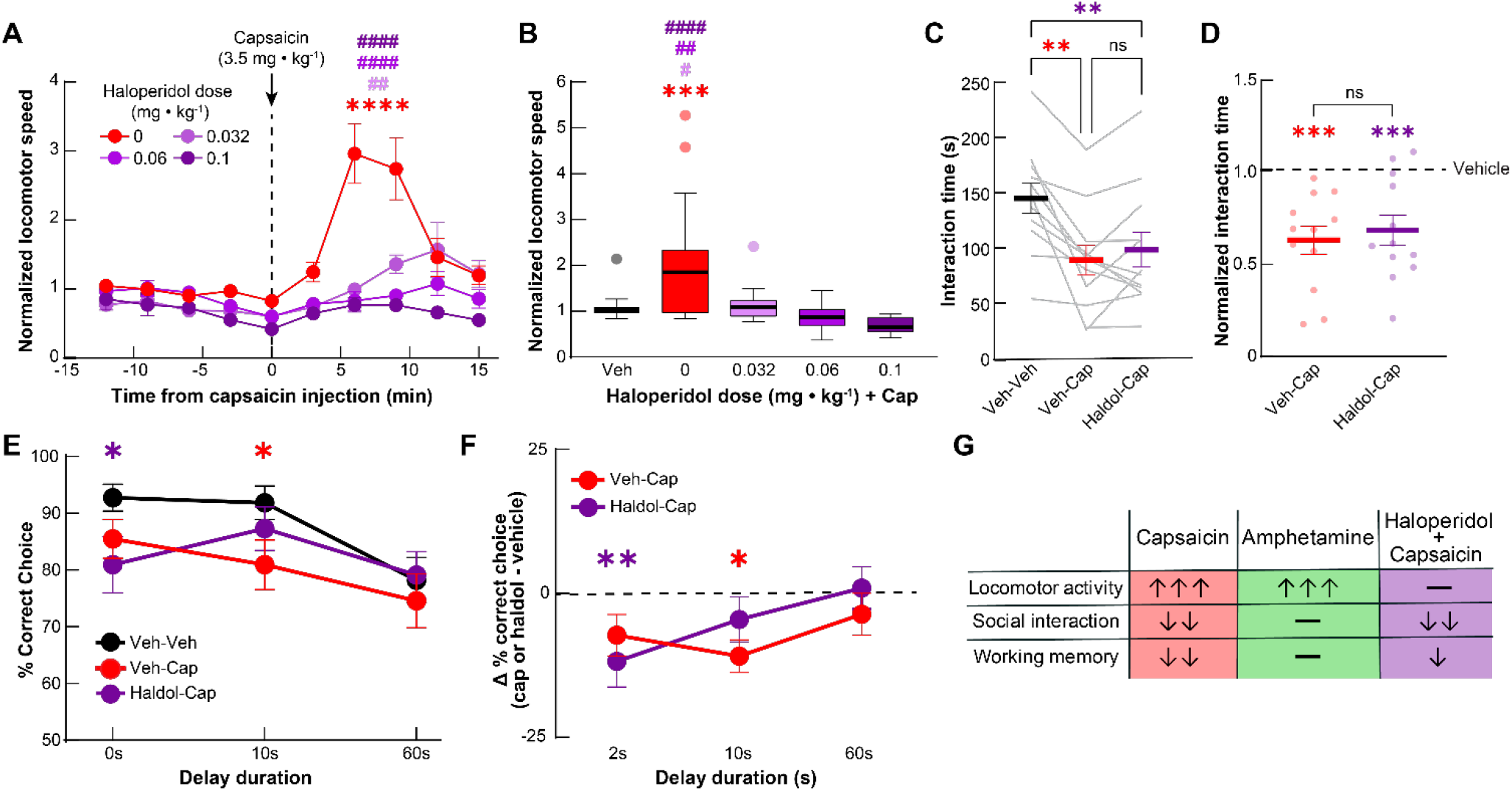
Haloperidol incompletely normalizes the effects of selective nigrostriatal dopamine excess. **A, B)** Pre-treating TRPV1 mice with the first-generation antipsychotic haloperidol suppressed the increase in locomotion caused by systemic capsaicin treatment (3.5 mg·kg^-1^). Data in **A** are mean ± s.e.m. normalized to values following vehicle treatment and plotted in 3-min time bins. Data in **B** are averaged during the first 15-min following capsaicin treatment, normalized to values following vehicle treatment (*N* = 26–29 TRPV1 mice). **C, D)** Haloperidol pre-treatment (1 mg·kg^-1^) failed to normalize the decrease in juvenile social interaction time in experimental mice caused by capsaicin treatment (3.5 mg·kg^-1^), **C**, or the change in social interaction time, **D**, between capsaicin and vehicle treatment (mean ± s.e.m; *N* = 12 TRPV1 mice). **E, F)** Mean ± s.e.m. percent correct choice, **E**, and change in percent choice from vehicle treatment following capsaicin treatment (3.5 mg·kg^-1^), **F**, in the DNMS T-maze task was reduced in TRPV1 mice and only partially rescued by haloperidol pre-treatment (*N* = 11 TRPV1 mice). **G)** Summary of behavioral results in the different experimental groups included in this study. Arrows indicate directionality and magnitude of effects. Holm-Sidak’s multiple comparison test compared to vehicle (\*\*\*\**P* < 0.0001, \*\*\**P* < 0.001) and vehicle + capsaicin (^####^*P* < 0.0001, ^##^*P* < 0.01 and ^#^*P* < 0.05) in **A, B**, or to vehicle-vehicle (\*\*\**P* < 0.001, \*\**P* < 0.01 and \**P* < 0.05) treatment in **C–F**).

Taken together, these data show that haloperidol normalized behavioral processes that are typically normalized by antipsychotic drugs in rodents (*i*.*e*., hyperlocomotion). However, haloperidol failed to normalize, or only partially normalized behavioral processes associated with the treatment-resistant symptoms of psychosis in patients (social interaction and working memory, respectively) (**Figure 6G**). Despite blocking striatally enriched D2Rs, haloperidol was only partially effective for the behavioral changes driven by a selective excess in DS dopamine.

## DISCUSSION

The adverse effects of cognitive and negative symptoms in psychosis are widely recognized, yet their underlying etiology is poorly understood. The prevailing view is that excess dopamine is not involved in these symptoms, since antipsychotic drugs that block dopamine receptors do not alleviate these symptoms, and drugs that increase dopamine do not reliably induce them. In the current study, we directly tested this idea in a manner that explicitly models the pathway-specific increase in dopamine observed in schizophrenia patients.

To do this, we adapted an approach that was previously used to activate all dopamine neurons (21). Specifically, we generated a virus to selectively express excitatory TRPV1 receptors in DS-projecting, SNc dopamine neurons of DAT^cre/+^ mice that lack the endogenous *Trpv1* gene. In contrast to the transgenic approach used in previous studies to selectively express TRPV1 (21, 22), our approach allowed us to specifically activate genetically defined neurons at the site of virus injection. This virus-mediated approach is widely applicable to manipulating circuits throughout the brain. We leveraged this regional and genetic selectivity to transiently activate DS-projecting SNc neurons (but not other dopamine neurons) using the TRPV1 agonist capsaicin and selectively induced dopamine release in the DS, but not mPFC. Capsaicin’s effects on behavior and dopamine release lasted approximately 15–20 min, consistent with earlier studies using capsaicin to activate all dopamine neurons (21). This transient duration is an advantage for experimental designs requiring a modest duration (minutes) of reversible activation, but a limitation for experiments requiring shorter or longer stimulations (subseconds or hours). Other advantages include the compatibility with traditional chemogenetic approaches (since capsaicin and DREADD ligands are distinct) and *in vivo* imaging approaches that are constrained by limited cranial space for implanting optogenetic fibers (18, 25, 26). A notable limitation of this approach is the requirement that the mice must be on a *Trpv1* knockout background, which requires at least two generations of crosses to generate experimental animals. The lack of endogenous TRPV1 expression also has the potential to confound behavioral phenotypes. Although here we used control mice that also lacked the *Trpv1* gene, this could particularly confound studies of behavioral processes such as sensation or pain that are influenced by TRPV1 (17).

Treating TRPV1 mice with capsaicin and C57BL6/J mice with amphetamine both induced hyperlocomotion. A drug’s ability to suppress amphetamine-driven locomotion in mice is commonly used to assess its antipsychotic potential in humans (23). Albeit circular, there is logic to this idea. Most antipsychotic drugs attenuate dopamine receptor signaling, and hyperlocomotion results from amphetamine-induced increases in striatal dopamine release (27). Because both amphetamine-driven locomotion in rodents and the positive symptoms of schizophrenia in patients respond to antipsychotic treatment, they are speculated to engage partly overlapping neural substrates. Although further experiments are necessary to confirm that capsaicin treatment induces behavioral changes associated with positive symptoms in TRPV1 mice, the fact that it induces haloperidol-responsive hyperlocomotion suggests it is engaging dopamine pathways related to the positive symptoms of psychosis.

Although capsaicin (3.5 mg·kg^-1^ in TRPV1 mice) and amphetamine (1 mg·kg^-1^ in C57BL6/J mice) treatments equivalently induced hyperlocomotion, only the former additionally perturbed social interaction and working memory. Social exploration and working memory are complex processes that rely on distributed brain areas (20, 28-30). The mPFC, in particular, is crucially involved in both processes (31-33), and the brains of schizophrenia patients exhibit an array of pathological changes in the prefrontal cortex that are thought to underlie cognitive and negative symptoms (34). In fact, the onset of these symptoms can precede that of the positive symptoms, further supporting distinct underlying etiologies (35, 36). However, patients with prodromal schizophrenia symptoms also have elevated dopamine signaling in the DS that correlates with neurocognitive dysfunction (37). Furthermore, the DS is both directly and indirectly connected to the brain structures most implicated in cognitive and social function. For instance, inhibiting DMS-projecting mPFC neurons during the delay period of a T-maze working memory task impairs performance, and DMS neurons are sequentially active during this delay period (13, 38). The DS is also connected through basal ganglia outputs to the thalamocortical connections thought to sustain persistent activity in the PFC during delay periods of working memory (20, 39). Striatal dysfunction has also been repeatedly implicated in social behavior deficits, particularly in the context of autism-spectrum disorders (40-42). Moreover, optogenetically activating nigrostriatal dopamine neurons was recently shown to reduce social preference and decrease social interaction in mice (40). Therefore, our own findings are consistent with previous studies implicating striatal function in social interaction and working memory. Our findings add to this understanding by distinguishing nigrostriatal from brain-wide dopamine excess (*i*.*e*., following amphetamine treatment), which did not disrupt social interaction or working memory. This distinction could reflect compensation due to dopamine release in brain regions outside of the striatum (*e*.*g*., in the PFC) following amphetamine treatment, or the brain-wide competition for cognitive resources between structures when dopamine release is asymmetrically driven (15). Intriguingly, a seemingly inverse relationship between striatal and cortical dopamine signaling has long been recognized in humans and in animal studies of schizophrenia, and this circuit-level interaction between the two structures could explain the diversity of psychosis symptoms (43-49). This interdependence and its implications for regional brain dysfunction further underscore the importance of addressing excess striatal dopamine in psychosis.

If nigrostriatal dopamine excess contributes to cognitive and negative symptoms, then why are antipsychotics that block D2 dopamine receptors largely ineffective for these symptoms? The simplest explanation is that these drugs fail to properly engage the neural substrates responsible for these symptoms (*e*.*g*., decreased dopamine, glutamate, or neuropathology in the PFC). While a lack of effects outside of the striatum undoubtedly contributes to their limited efficacy, even within the striatum, antipsychotics do not fully address the consequences of dopamine excess. Approximately half of the neurons in the striatum express D1 rather than D2Rs. Because antipsychotics do not principally target them, striatal D1Rs could contribute to the treatment-resistant symptoms. Consistent with this idea, we recently found that clozapine, a highly efficacious antipsychotic drug with some pro-cognitive effects, preferentially normalized activity in D1R-expressing spiny projection neurons (SPNs) in the DMS under hyperdopaminergic conditions (50, 51). By contrast, haloperidol affected both D1- and D2-SPN activity following amphetamine treatment (50). These data potentially explain why haloperidol partially rescued working memory in capsaicin-treated TRPV1 mice, and suggests that drugs that preferentially normalize D1-SPN activity may have greater therapeutic advantages over drugs like haloperidol that principally block D2Rs (50).

In summary, our results show that excess dopamine, when restricted to the dorsal striatum, alters behavioral processes linked to the positive, negative, and cognitive symptom subclasses of psychosis. Non-selectively driving dopamine release did not disrupt the processes associated with negative and cognitive symptoms, and haloperidol treatment only partially normalized the deficits in these behaviors in our model. Our results implicate nigrostriatal hyperdopaminegia in the etiology of psychosis symptoms that do not respond to current treatments, but other brain areas are also undoubtedly involved, particularly the PFC. Within the context of our earlier studies, treatments targeted to striatal D1-, rather than D2-SPNs may have therapeutic benefits over classical antipsychotic drugs. Further studies are necessary to unravel the intricacies of nigrostriatal dopamine excess in the behavioral processes examined here. In particular, the DS is a large, heterogeneous brain structure encompassing several subregions that contain multiple cell-types innervated by multiple dopamine cell sub-types. A better understanding of these intricacies holds the promise to develop more targeted therapies that better address the diverse symptoms of psychosis.

## ACKNOWLEDGEMENTS AND DISCLOSURES

This work was supported by the National Institutes of Mental Health: K01 MH11313201 (JGP) and 2T32 MH067564 (NAM), Neurological Disorders and Stroke R01 NS122840 (JGP), Drug Abuse P30DA048736 (LSZ) and Diabetes and Digestive and Kidney Diseases R01 DK128477 (LRB). We thank Dr. Ben Arenkiel for providing the cDNA plasmid for *Trpv1*. The authors have no financial disclosures to declare.

## SUPPLEMENTARY METHODS

### Viruses

We synthesized a polylinker containing a 5’ SpeI and 3’ PacI site (5’CTAGCACTATGAGCTCTTAATTAAG 3’ and 5’CTAGCTTAATTAAGAGCTCACTAGTG3’) and inserted it into the NheI site of pAAV-hSyn-FLEX-mCherry (Addgene #50459). We PCR amplified rat *Trpv1* (forward primer: 5’GATATCTTAATTAAATGGAACAACGGGCTAGCTTA3’ and reverse primer: 5’GATATCACTAGTTTTCTCCCCTGGGACCATGGA3’; IDT) and inserted the amplified construct in-frame with mCherry with PacI and SpeI (NEB) using restriction digest cloning. We sent the resulting plasmid to Virovek for virus production (AAV2/5-FLEX-TRPV1-mCherry). Virovek also produced AAV2/9-CAG-dLight1.3b directly from the Addgene plasmid (#125560). We purchased a viral preparation of AAV2/5-Syn-DIO-mCherry directly from Addgene (#50459).

### Surgical Procedures

For virus injections, we anesthetized mice with isoflurane (2% in O_2_) and stereotaxically injected virus at a rate of 250 nL·min^-1^ into each region of interest using a microsyringe with a 33-gauge beveled tip needle. All anteroposterior (AP), mediolateral (ML) coordinates are reported from bregma and dorsoventral (DV) coordinates are reported from bregma or dura as specified. For all virus injections, we went 0.5-mm (DV) past the injection target and then withdrew the syringe and needle back to the target for the injection. After each injection, we left the syringe in place for 5 min, withdrew the syringe 0.1 mm, and then waited 5 additional min before slowly with-drawing the syringe to minimize viral spread. To express TRPV1-mCherry or control mCherry in SNc dopamine neurons, we injected 500 nL of AAV2/5-Syn-FLEX-TRPV1-mCherry (8.4 × 10^12^ GC/mL) or AAV2/5-hSyn-DIO-mCherry (1.7 × 10^12^ GC/mL) bilaterally into the SNc (AP: [bregma to lambda distance]/4.21 × -3.5 mm, ML: ±1.5 mm, and DV: -4.0 mm from bregma). To express dLight1.3b in the DMS and mPFC, we injected 500 nL of AAV2/9-CAG-dlight1.3b (1.53 × 10^12^ GC/mL) unilaterally into the mPFC (AP: [bregma to lambda]/4.21 × +1.9 mm, ML= -0.25 mm and DV= -2.1 mm from dura) and DMS (AP: +0.5 mm, ML: -1.5 mm and DV: -2.7 mm from dura).

For mice only receiving virus injections, we then sutured the scalp, injected analgesic subcutaneously (Buprenorphine SR; 1 mg·kg^-1^) and waited 3–4 weeks for the mice to recover and for viral expression before beginning behavioral experiments. For mice undergoing fiber-optic implantation for photometry, in the same surgery we applied 4 stainless steel screws at arbitrary locations in the skull (AMS120/1; Antrin) and implanted fiberoptic cannulas (MFC_400/430-0.48_4mm; Doric) into the same AAV injection coordinates in the mPFC and DMS. After placing the fiberoptic cannulae, we applied Metabond (Parkell) to the skull and used dental acrylic (Coltene) to fix the full assembly along with a custom stainless-steel head-plate (Laser Alliance) for head-fixing mice during attachment and release of photometry patch cords. We injected analgesic and allowed the mice to recover for 3–4 weeks before recording dlight1.3b fluorescence.

### Fiber Photometry

We tethered each mouse’s implanted fiberoptic cannulas to low-autofluorescence patch cords (MFP_400/430/1100-0.57_2m_LAF; Doric). We used a controller and LED sources (Thorlabs) with miniature filter cubes (FMC6_AE(400-410)_E1(460-490)_F1(500-540)_E2(550-580)_F2(600-680)_S; Doric) to deliver ∼30–70 μW of blue (470 nm) and UV (405 nm) light modulated at 211 and 511 Hz, respectively. We recorded dLight1.3b green fluorescence signals during blue and UV (isosbestic) light absorption through the same patch cords coupled to femtowatt photoreceivers (2151; Newport). We used a commercial system and software (RZ5P and Synapse; TDT) to sample (at 1.0173 kHz), de-modulate, and lock-in amplify these digital signals.

We habituated mice to recording by tethering the patch cord to their implanted fiberoptic cannulas and allowing them to explore an open field arena in two, 1-h sessions on consecutive days. For fiber photometry recordings, we recorded each mouse’s baseline dLight1.3b activity in the mPFC and DMS for 15 min, subcutaneously injected vehicle, recorded for an additional 15 min, and then subcutaneously injected capsaicin or amphetamine (10 mg·kg^-1^ each) and recorded dLight1.3b activity for 30 or 60 min, respectively. We then used custom MATLAB scripts to compute normalized dLight1.3b fluorescence traces (% Δ*F/F*) by using a polynomial fit to adjust the isosbestic (405 nm) fluorescence signal trace to that of the blue-light (470 nm) fluorescence trace and then subtracting it from the 470-nm fluorescence trace and multiplying the resulting trace by 100. We ignored the baseline recording period and removed spurious outlier points from the Δ*F/F* traces following vehicle and drug treatment that were greater or less than ± 3 standard deviations (SD) of the mean value for the trace during the given treatment. We then applied a 1-s median filter to temporally smooth the trace and averaged the trace to down sample its temporal resolution to 1 Hz. Next, we computed the area-under-the-curve (AUC) of the trace in 1-min time bins. We calculated the average AUC of the last 10, 1-min bins of the vehicle recording and subtracted this value from the AUC observed in every 1-min time bin following vehicle and drug treatment (**Figure 2E–H**). We then identified the 1-min time bin in which the normalized AUC was maximal following drug treatment and computed the average AUC for that bin and the 4, 1-min bins before and 5, 1-min bins after (average of 10-min about the maximum; **Figure 2I, J**). To eliminate potential false negative results due to the inability to detect dopamine signaling in the mPFC, we excluded mice where the average AUC of the 10-min bins around the maximum following amphetamine treatment did not exceed 2 SD of the mean AUC (across mice) of the 10, 1-min bins at the end of the vehicle treatment recording. This criterion resulted in 5/18 mice being excluded from the mPFC analysis due to the lack of a significant increase in dLight1.3b fluorescence following amphetamine treatment (**Figure 2F, H, J**). Using the same criterion, all mice had a significant increase in DMS dLight1.3b fluorescence following amphetamine treatment, so we excluded no mice from our analysis of amphetamine or capsaicin effects on DMS dopamine signaling (**Figure 2E, G, I**).

### Open Field Locomotor Activity

We habituated mice in 1-h sessions to a clear acrylic circular open field arena (30.48-cm diameter; Tap plastics) for two days. During these habituation sessions, we also administered one subcutaneous injection of vehicle (3.33% Tween-80 in PBS or saline) and one subcutaneous injection of capsaicin (10 mg·kg^-1^) or amphetamine (2.5 mg·kg^-1^) to habituate them to drug injection. On testing days, we habituated mice to the open field for 20 min, injected vehicle, recorded locomotion for 30 min or 1 h, injected capsaicin or amphetamine at varying doses (one per day) and recorded locomotion for 30 min or 1 h (**Figure 3B, 5A**). For haloperidol experiments, we pretreated mice with vehicle or varying doses of haloperidol, recorded locomotion for 30 min, injected capsaicin (3.5 mg·kg^-1^) and again recorded locomotion for 30 min (**Figure 6A**). To analyze locomotor speed, we used a video camera and software (DMK 22BUC03 and IC Capture 2.4 [The Imaging Source]) with a varifocal lens (T3Z2910CS; Computar) to record 30-Hz videos of freely moving mouse behavior. We used custom software written in ImageJ to track each mouse’s position in an open field arena (https://bahanonu.github.io/ciatah/) (18).

### Spatial Working Memory

We individually housed mice and gradually food restricted them to 85% of their *ad libitum* body weight. Once at 85% body weight, we habituated mice to an automated T-maze (MazeEngineers) for 2 days. On day 1 of habituation, we placed mice in the maze and allowed them to freely explore and forage for 45 mg chocolate-flavored pellets (Bio-Serv) sporadically placed throughout the maze for 30 min. On day 2 of habituation, mice explored the maze for 30 min and received a food reward delivery each time they entered either of the maze goal arms. On the subsequent 2 days, we pre-trained mice in sessions consisting of 10 alternating (left vs. right) forced trials from the center holding area to a baited goal arm and then back to the center holding area. The day after pretraining, we trained mice in a delayed non-match to sample (DNMS) task in which we interleaved forced and choice trials (20). During the choice trial, mice were required to choose the arm opposite to the immediately preceding forced trial in order to obtain a food reward. We pseudo-randomly chose the ordering of left and right forced trials.

Once mice had at least three consecutive days with ≥70% correct choice trials, we proceeded with pharmacological testing. In each session, we injected mice with vehicle, capsaicin (3.5 mg·kg^-1^), or amphetamine (1 mg·kg^-1^), waited 2 min (capsaicin) or 4 min (amphetamine), and tested their DNMS performance with a delay period of 2, 10, or 60 s between forced and choice trials on sequential days (**Figure 4E–G, 5F–G**). We administered successive capsaicin injections every 15 min throughout each session to follow the time course of optimal drug effect (**Figure 3B**). For haloperidol experiments, we pretreated mice with either vehicle or haloperidol (0.1 mg·kg^-1^), waited 10 min, then injected capsaicin (3.5 mg·kg^-1^), waited 2 min, and tested their performance with the same 2, 10, and 60 s delays while administering capsaicin every 15-min until task completion (**Figure 6E–G**)

### Histology

After all photometry and behavioral experiments, we euthanized and intracardially perfused mice with PBS followed by a 4% solution of paraformaldehyde in PBS. For the mice used for Fos quantification, we injected vehicle or capsaicin (10 mg·kg^-1^) 75 min before euthanasia. After perfusion, we extracted the brains and placed them in 4% paraformaldehyde for 1–3 days. We sliced 50–100-μM-thick coronal sections from the fixed-brain tissue using a vibratome (Leica VT1000s). For immunostaining, we used α-Tyrosine Hydroxylase (1:500; Aves TYH) or α-Fos (1:1000; Cell Signaling #2250) primary antibodies in combination with α-RFP (1:1000; Rockland #200-301-379) and Alexa 488 with 594 conjugated secondary antibodies (1:500; Jackson Immunoresearch #703-546-155 or #711-546-152 with #715-586-150, respectively) to amplify and label mCherry-expressing SNc neurons in alternating midbrain sections (**Figure 1**). For dlight1.3b analysis in DMS and mPFC, we used an anti-GFP antibody (1:1000, Invitrogen #A11122) and Alexa 488 conjugated secondary antibody (**Figure 2B**; 1:500, Jackson Immunoresearch #711-546-152). We then mounted the sections with DAPI-containing mounting media (Southern Biotech #0100-20) and imaged fluorescence using a two-photon (Bruker) or wide-field, slide-scanning fluorescence microscope (Keyence BZ-X800 or Leica THUNDER) with 10×, 16×, or 20× objectives. We used ImageJ and opensource plugins to process all data and quantify Fos or TH colocalization with mCherry in SNc neurons (https://imagej.nih.gov/ij/plugins/cell-counter.html) (**Figure 1C, E**).

## SUPPLEMENTARY TABLE

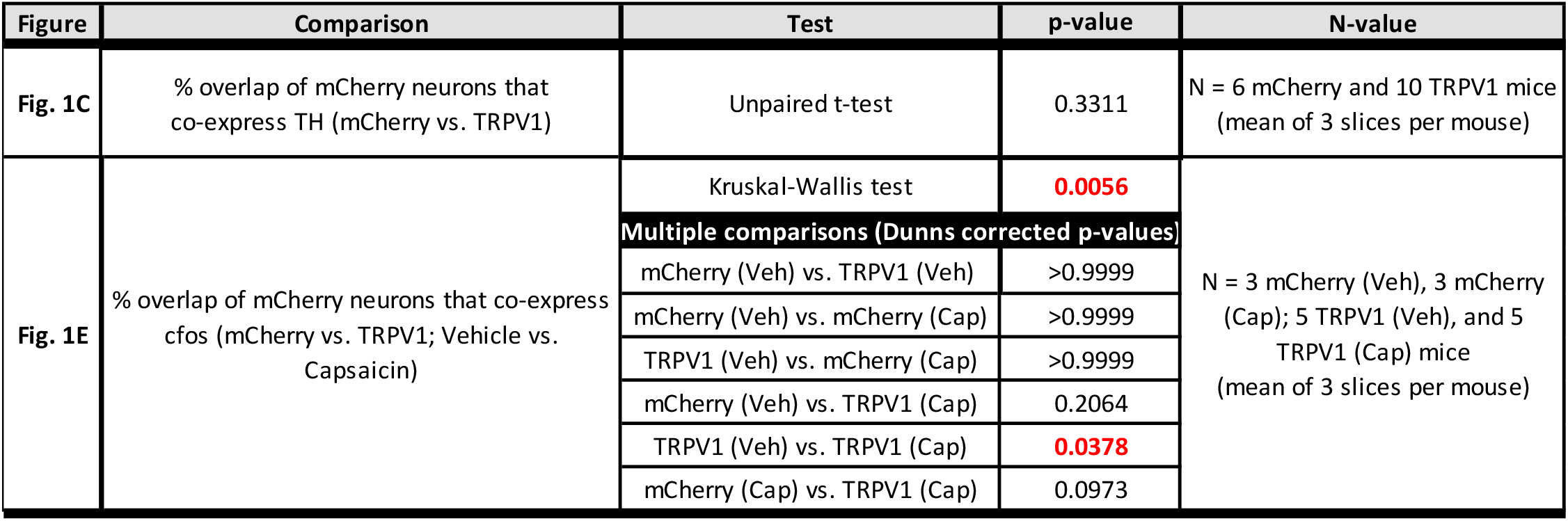

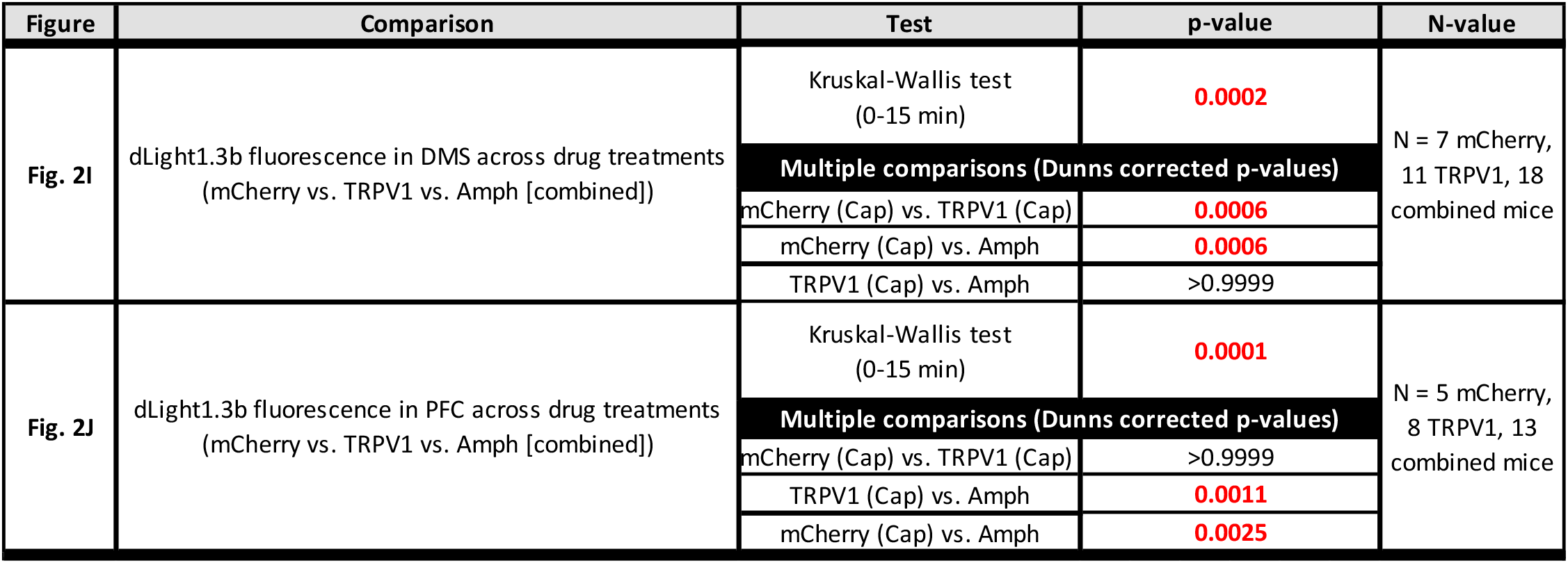

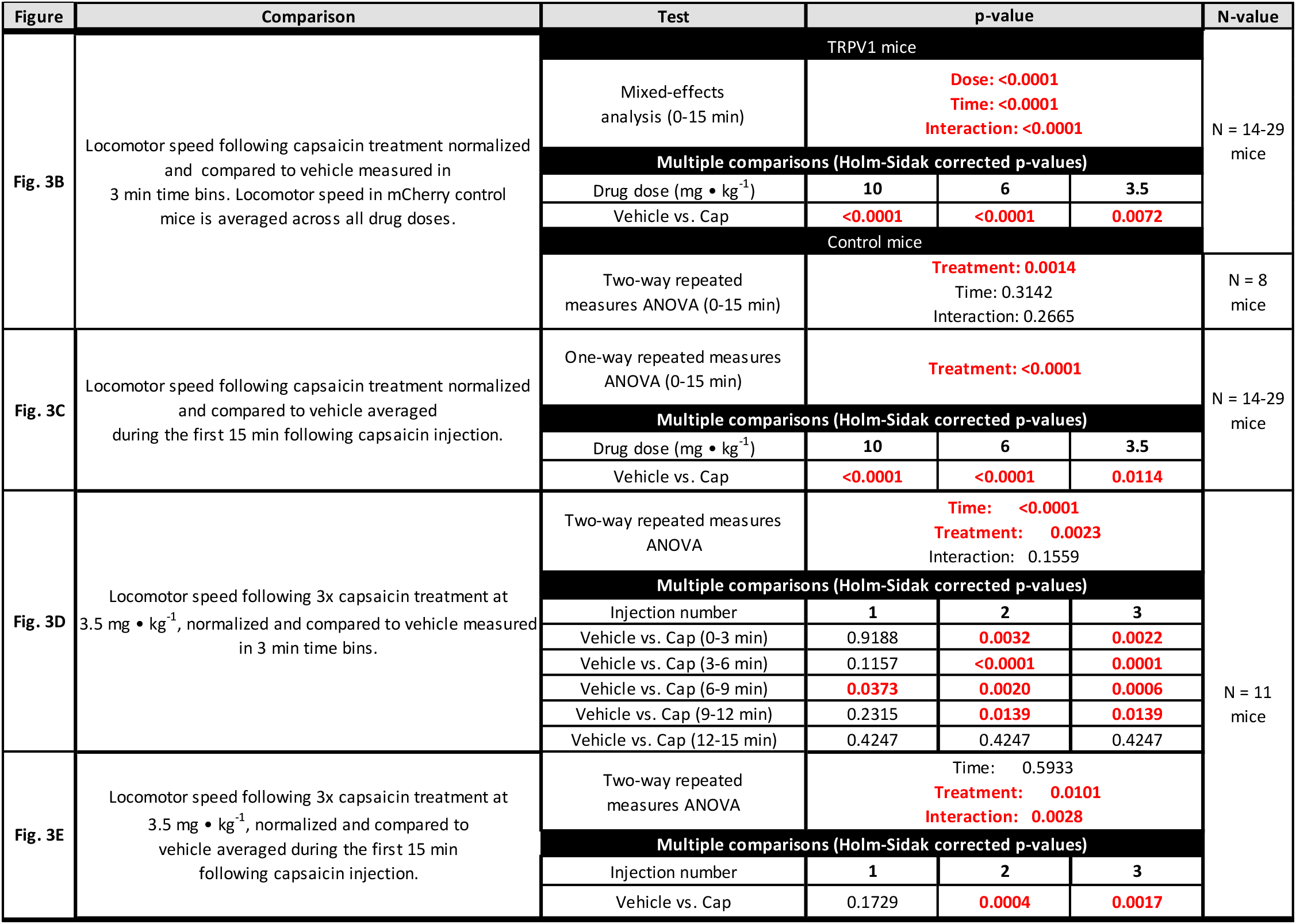

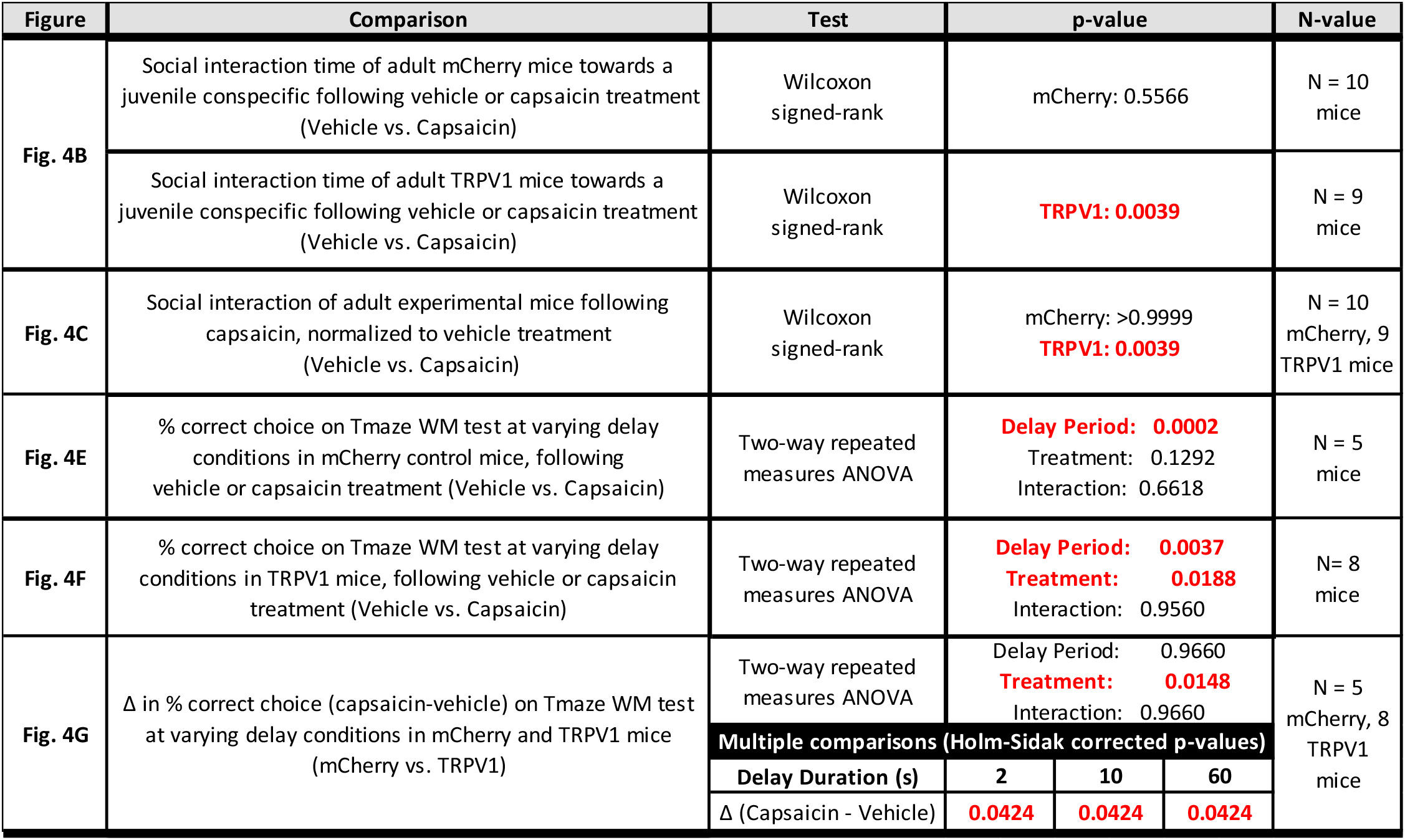

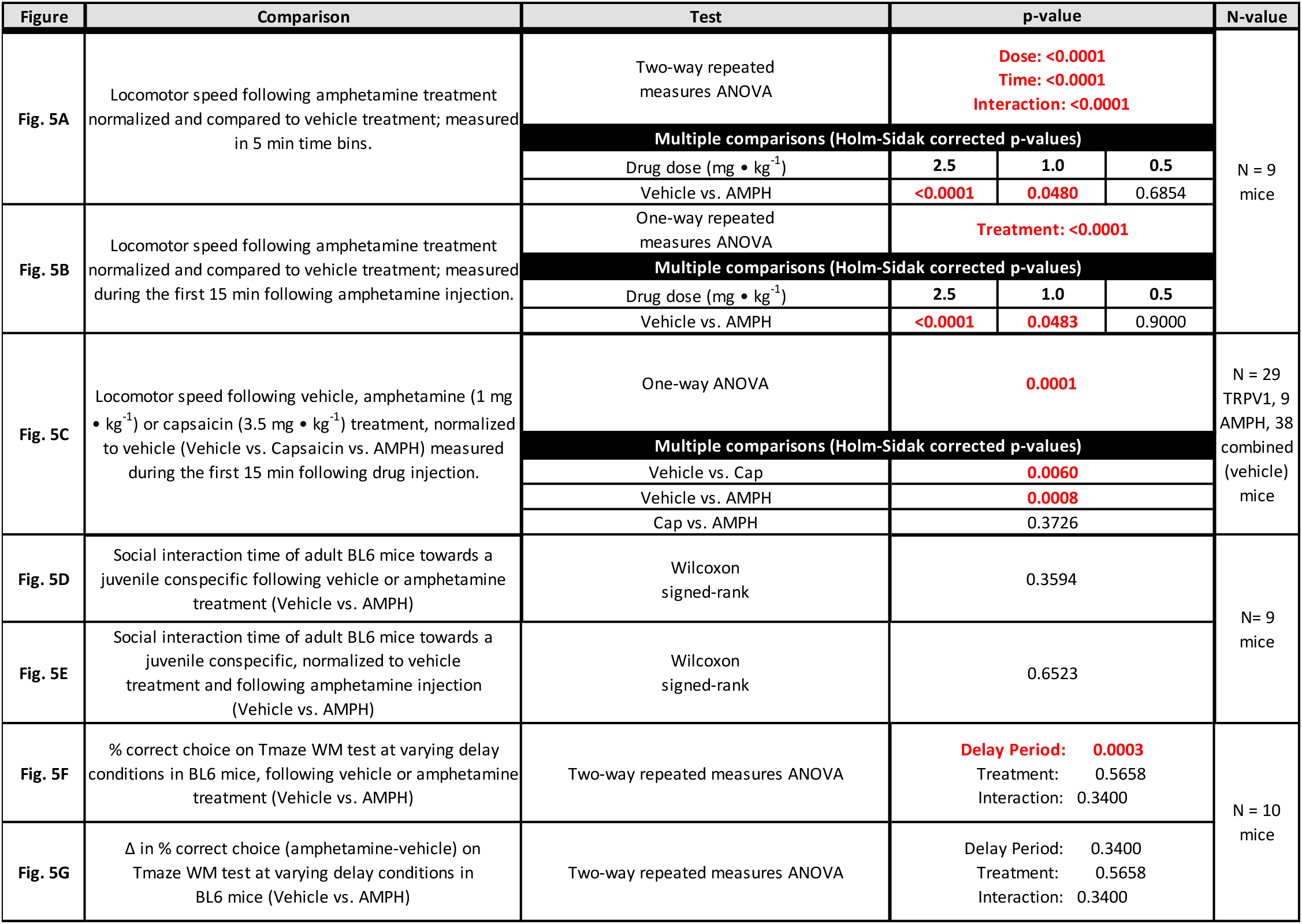

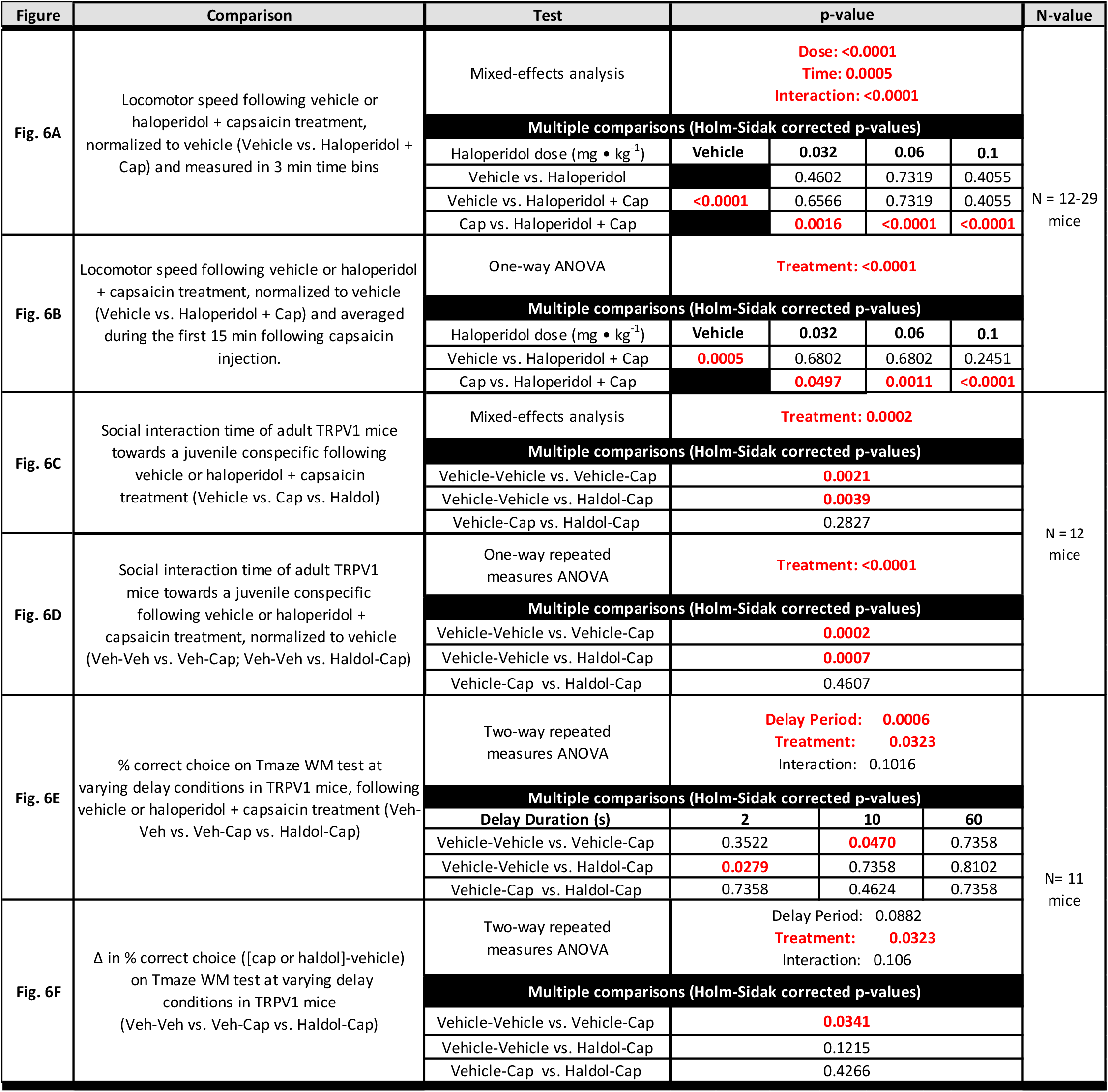

## REFERENCES

1. Huhn M, Nikolakopoulou A, Schneider-Thoma J, Krause M, Samara M, Peter N, et al. (2019): Comparative efficacy and tolerability of 32 oral antipsychotics for the acute treatment of adults with multi-episode schizophrenia: a systematic review and network meta-analysis. Lancet. 394:939–951.

2. Harvey PD, Strassnig M (2012): Predicting the severity of everyday functional disability in people with schizophrenia: cognitive deficits, functional capacity, symptoms, and health status. World Psychiatry. 11:73–79.

3. Seeman P, Lee T, Chau-Wong M, Wong K (1976): Antipsychotic drug doses and neuroleptic/dopamine receptors. Nature. 261:717–719.

4. Creese I, Burt DR, Snyder SH (1976): Dopamine receptor binding predicts clinical and pharmacological potencies of antischizophrenic drugs. Science. 192:481–483.

5. Abi-Dargham A, Rodenhiser J, Printz D, Zea-Ponce Y, Gil R, Kegeles LS, et al. (2000): Increased baseline occupancy of D2 receptors by dopamine in schizophrenia. Proc Natl Acad Sci U S A. 97:8104–8109.

6. Hurd YL, Suzuki M, Sedvall GC (2001): D1 and D2 dopamine receptor mRNA expression in whole hemisphere sections of the human brain. J Chem Neuroanat. 22:127–137.

7. Davis KL, Kahn RS, Ko G, Davidson M (1991): Dopamine in schizophrenia: a review and reconceptualization. Am J Psychiatry. 148:1474–1486.

8. Bell DS (1973): The experimental reproduction of amphetamine psychosis. Arch Gen Psychiatry. 29:35–40.

9. Barch DM, Carter CS (2005): Amphetamine improves cognitive function in medicated individuals with schizophrenia and in healthy volunteers. Schizophr Res. 77:43–58.

10. Missale C, Nash SR, Robinson SW, Jaber M, Caron MG (1998): Dopamine receptors: from structure to function. Physiol Rev. 78:189–225.

11. Slifstein M, van de Giessen E, Van Snellenberg J, Thompson JL, Narendran R, Gil R, et al. (2015): Deficits in prefrontal cortical and extrastriatal dopamine release in schizophrenia: a positron emission tomographic functional magnetic resonance imaging study. JAMA Psychiatry. 72:316–324.

12. McCutcheon RA, Abi-Dargham A, Howes OD (2019): Schizophrenia, Dopamine and the Striatum: From Biology to Symptoms. Trends Neurosci. 42:205–220.

13. Akhlaghpour H, Wiskerke J, Choi JY, Taliaferro JP, Au J, Witten IB (2016): Dissociated sequential activity and stimulus encoding in the dorsomedial striatum during spatial working memory. eLife. 5.

14. Graybiel AM (1997): The basal ganglia and cognitive pattern generators. Schizophr Bull. 23:459–469.

15. Poldrack RA, Packard MG (2003): Competition among multiple memory systems: converging evidence from animal and human brain studies. Neuropsychologia. 41:245–251.

16. Bäckman CM, Malik N, Zhang Y, Shan L, Grinberg A, Hoffer BJ, et al. (2006): Characterization of a mouse strain expressing Cre recombinase from the 3’ untranslated region of the dopamine transporter locus. Genesis. 44:383–390.

17. Caterina MJ, Leffler A, Malmberg AB, Martin WJ, Trafton J, Petersen-Zeitz KR, et al. (2000): Impaired nociception and pain sensation in mice lacking the capsaicin receptor. Science. 288:306–313.

18. Parker JG, Marshall JD, Ahanonu B, Wu YW, Kim TH, Grewe BF, et al. (2018): Diametric neural ensemble dynamics in parkinsonian and dyskinetic states. Nature. 557:177–182.

19. Fernandez Espejo E (2003): Prefrontocortical dopamine loss in rats delays long-term extinction of contextual conditioned fear, and reduces social interaction without affecting short-term social interaction memory. Neuropsychopharmacology. 28:490–498.

20. Bolkan SS, Stujenske JM, Parnaudeau S, Spellman TJ, Rauffenbart C, Abbas AI, et al. (2017): Thalamic projections sustain prefrontal activity during working memory maintenance. Nat Neurosci. 20:987–996.

21. Guler AD, Rainwater A, Parker JG, Jones GL, Argilli E, Arenkiel BR, et al. (2012): Transient activation of specific neurons in mice by selective expression of the capsaicin receptor. Nature communications. 3:746.

22. Dietrich MO, Zimmer MR, Bober J, Horvath TL (2015): Hypothalamic Agrp neurons drive stereotypic behaviors beyond feeding. Cell. 160:1222–1232.

23. van den Buuse M (2010): Modeling the positive symptoms of schizophrenia in genetically modified mice: pharmacology and methodology aspects. Schizophr Bull. 36:246–270.

24. Moghaddam B, Bunney BS (1989): Differential effect of cocaine on extracellular dopamine levels in rat medial prefrontal cortex and nucleus accumbens: comparison to amphetamine. Synapse. 4:156–161.

25. Roth BL (2016): DREADDs for Neuroscientists. Neuron. 89:683–694.

26. Beutler LR, Chen Y, Ahn JS, Lin YC, Essner RA, Knight ZA (2017): Dynamics of Gut-Brain Communication Underlying Hunger. Neuron. 96:461-475.e465.

27. Kelley AE, Gauthier AM, Lang CG (1989): Amphetamine microinjections into distinct striatal subregions cause dissociable effects on motor and ingestive behavior. Behav Brain Res. 35:27–39.

28. Kim Y, Venkataraju KU, Pradhan K, Mende C, Taranda J, Turaga SC, et al. (2015): Mapping social behavior-induced brain activation at cellular resolution in the mouse. Cell reports. 10:292–305.

29. Guo ZV, Inagaki HK, Daie K, Druckmann S, Gerfen CR, Svoboda K (2017): Maintenance of persistent activity in a frontal thalamocortical loop. Nature. 545:181–186.

30. McNab F, Klingberg T (2008): Prefrontal cortex and basal ganglia control access to working memory. Nat Neurosci. 11:103–107.

31. Yizhar O, Levy DR (2021): The social dilemma: prefrontal control of mammalian sociability. Curr Opin Neurobiol. 68:67–75.

32. Funahashi S, Chafee MV, Goldman-Rakic PS (1993): Prefrontal neuronal activity in rhesus monkeys performing a delayed anti-saccade task. Nature. 365:753–756.

33. Kim IH, Kim N, Kim S, Toda K, Catavero CM, Courtland JL, et al. (2020): Dysregulation of the Synaptic Cytoskeleton in the PFC Drives Neural Circuit Pathology, Leading to Social Dysfunction. Cell reports. 32:107965.

34. Lewis DA (2012): Cortical circuit dysfunction and cognitive deficits in schizophrenia--implications for preemptive interventions. Eur J Neurosci. 35:1871–1878.

35. Piskulic D, Addington J, Cadenhead KS, Cannon TD, Cornblatt BA, Heinssen R, et al. (2012): Negative symptoms in individuals at clinical high risk of psychosis. Psychiatry Res. 196:220–224.

36. Bora E, Lin A, Wood SJ, Yung AR, McGorry PD, Pantelis C (2014): Cognitive deficits in youth with familial and clinical high risk to psychosis: a systematic review and meta-analysis. Acta Psychiatr Scand. 130:1–15.

37. Howes OD, Montgomery AJ, Asselin MC, Murray RM, Valli I, Tabraham P, et al. (2009): Elevated striatal dopamine function linked to prodromal signs of schizophrenia. Arch Gen Psychiatry. 66:13–20.

38. Chernysheva M, Sych Y, Fomins A, Alatorre Warren JL, Lewis C, Capdevila LS, et al. (2021): Striatum-projecting prefrontal cortex neurons support working memory maintenance. bioRxiv.2021.2012.2003.471159.

39. McElvain LE, Chen Y, Moore JD, Brigidi GS, Bloodgood BL, Lim BK, et al. (2021): Specific populations of basal ganglia output neurons target distinct brain stem areas while collateralizing throughout the diencephalon. Neuron. 109:1721-1738.e1724.

40. Lee Y, Kim H, Kim JE, Park JY, Choi J, Lee JE, et al. (2018): Excessive D1 Dopamine Receptor Activation in the Dorsal Striatum Promotes Autistic-Like Behaviors. Mol Neurobiol. 55:5658–5671.

41. Kim H, Lee Y, Park JY, Kim JE, Kim TK, Choi J, et al. (2017): Loss of Adenylyl Cyclase Type-5 in the Dorsal Striatum Produces Autistic-Like Behaviors. Mol Neurobiol. 54:7994–8008.

42. Peça J, Feliciano C, Ting JT, Wang W, Wells MF, Venkatraman TN, et al. (2011): Shank3 mutant mice display autistic-like behaviours and striatal dysfunction. Nature. 472:437–442.

43. Krabbe S, Duda J, Schiemann J, Poetschke C, Schneider G, Kandel ER, et al. (2015): Increased dopamine D2 receptor activity in the striatum alters the firing pattern of dopamine neurons in the ventral tegmental area. Proceedings of the National Academy of Sciences. 112:E1498.

44. Kellendonk C, Simpson EH, Polan HJ, Malleret G, Vronskaya S, Winiger V, et al. (2006): Transient and selective overexpression of dopamine D2 receptors in the striatum causes persistent abnormalities in prefrontal cortex functioning. Neuron. 49:603–615.

45. Wilkinson LS (1997): The nature of interactions involving prefrontal and striatal dopamine systems. Journal of psychopharmacology (Oxford, England). 11:143–150.

46. Pycock CJ, Kerwin RW, Carter CJ (1980): Effect of lesion of cortical dopamine terminals on subcortical dopamine receptors in rats. Nature. 286:74–77.

47. Li YC, Kellendonk C, Simpson EH, Kandel ER, Gao WJ (2011): D2 receptor overexpression in the striatum leads to a deficit in inhibitory transmission and dopamine sensitivity in mouse prefrontal cortex. Proc Natl Acad Sci U S A. 108:12107–12112.

48. Kim IH, Rossi MA, Aryal DK, Racz B, Kim N, Uezu A, et al. (2015): Spine pruning drives antipsychotic-sensitive locomotion via circuit control of striatal dopamine. Nat Neurosci. 18:883–891.

49. Casado-Sainz A, Gudmundsen F, Baerentzen SL, Lange D, Ringsted A, Martinez-Tejada I, et al. (2022): Dorsal striatal dopamine induces fronto-cortical hypoactivity and attenuates anxiety and compulsive behaviors in rats. Neuropsychopharmacology. 47:454–464.

50. Yun S, Yang B, Martin M, Yeh N-H, Contractor A, Parker J (2021): Modulating D1 rather than D2 receptor-expressing spiny-projection neurons corresponds to optimal antipsychotic effect. bioRxiv.

51. Lee MA, Thompson PA, Meltzer HY (1994): Effects of clozapine on cognitive function in schizophrenia. J Clin Psychiatry. 55 Suppl B:82–87.

